# Pharmacological efficacy of a synergistic composition based on activators of cAMP accumulation as a wound healing agent

**DOI:** 10.1101/734814

**Authors:** Artur Martynov, Tatyana Bomko, Tatyana Nosalskaya, Boris Farber, Ostap Brek

## Abstract

**Introduction:** Wound-healing dipyridamole- and papaverine-based aerosols (D1/D2) as activators of the accumulation of cyclic adenosine monophosphate are promising drugs that can accelerate wound healing in wound processes of various origins.

**Materials and methods:** 128 rats were used in the study, including 38 in a pharmacological experiment on a model of stencil wounds and 90 in an experiment that studied the effect of spray on the number of CD34 cells in the blood of rats with chemically induced immunodeficiency. Immunodeficiency was caused by the fivefold administration of cyclophosphamide and prednisone. The expression level of CD34 was determined using flow cytofluorimeter.

**Results and discussion:** Topical application of D1/D2 aerosol samples on the skin of rats contributed to a statistically significant acceleration of regeneration processes. In terms of the appearance of granulations and epithelialization of wounds, D1/D2 aerosols were superior to dexpanthenol ointment. The maximum effect from the use of D1/D2 was observed on the 60th day, and restoration of the physiological level of pluripotent cells was observed as early as on the 10th day after the start of spray application. By accelerating wound healing, dipyridamole with papaverine probably stimulate the division of stem cells at the periphery without enhancing bone marrow function.

## Introduction

One of the main problems in the effective treatment of severe complications of type 2 diabetes mellitus is diabetic foot syndrome. Over time, the loss of sensitivity of the lower limbs at the late stages of diabetic neuropathy leads to a foot injury and the appearance of diabetic ulcers. These ulcers do not heal for a very long time; they become contaminated by bacteria, and in the end, would require surgical intervention [1]. As of today, there is no effective remedy that could heal such trophic wounds. Experiments proved the ability of endogenous stem cells to significantly accelerate the wound healing process, particularly in diabetic patients (rat model) [2]. Another study has demonstrated that an excessive amount of cAMP in a wound can stimulate the migration of stem cells to the wound area and their rapid differentiation [3,4].

As an agent for the treatment of trophic ulcers, we have developed a pharmaceutical composition in the form of an aerosol based on a combination of dipyridamole, papaverine and ascorbic acid. Papaverine and dipyridamole are universal phosphodiesterase inhibitors [5–7] and inducers of cAMP accumulation. Ascorbic acid represents an indirect adenylate cyclase activator, and also, an inducer of cAMP accumulation and an important component of the tissue regeneration process in wounds [8]. Using an animal model of Charcot-Marie-Tooth human disorder, was showed that ascorbic acid (AA) represses PMP22 gene expression by acting on intracellular cAMP concentrations. In this work, authors present kinetics data on the inhibitory effect of AA upon adenylate cyclase activity [9].

cAMP and cGMP are intracellular second messengers involved in the transduction of various physiologic stimuli and regulation of multiple physiological processes, including vascular resistance, cardiac output, visceral motility, immune response, inflammation, neuroplasticity, vision, and reproduction. Intracellular levels of these cyclic nucleotide second messengers are regulated predominantly by the complex superfamily of cyclic nucleotide phosphodiesterase (PDE) enzymes. Cyclic nucleotide phosphodiesterases (PDEs) comprise a superfamily of metallophospho hydrolases that specifically cleave the 3′, 5′-cyclic phosphate moiety of cAMP and/or cGMP to produce the corresponding 5′-nucleotide. PDEs are critical determinants for modulation of cellular levels of cAMP and/or cGMP by many stimuli. Thus, the ubiquitously present PDEs play a pivotal role in regulating cell signalling via the breakdown of cAMP and cGMP. PDE inhibitors are therapeutic agents which target PDE isoenzymes and inhibit the metabolism of the secondary messengers (cAMP, cGMP) thus, prolonging the biological effect determined by the type of cell involved [10,11].

It has also been shown that dipyridamole is able to stimulate the division of stem cells, which are the precursors of neurons [12]. Dipyridamole effectively stimulates capillary growth in the myocardium of rats [13]. Dipyridamole has been found to be able to potentiate the action of acyclic nucleotides in the treatment of herpes viruses, probably through its immunomodulating interferonogenic effect [14–16].

Papaverine also has a number of additional immunotropic properties that have not been previously used in pharmacological practice. For example, papaverine has antiviral properties against a broad spectrum of viruses, due to both its direct antiviral effect and its ability to stimulate the production of endogenous interferons [17–19].

The goal of this study is to determine the ability of the combination of two phosphodiesterase inhibitors, papaverine and dipyridamole in combination with ascorbic acid, to accelerate the healing of uninfected wounds in an experiment.

The contradictory effects of the composition for the individual components the various wounds healing process were previously shown.

It was also previously shown that papaverine is able to accelerate the healing of anastomoses in rats, but slightly (about 15%), although statistically significantly [20]

Dipyridamole, on the contrary, inhibits wound healing by 25% against control [21]

The preservative 0.05% chlorhexidine has no effect on uninfected wounds, although it slightly accelerates the healing the infected wounds due to antiseptic properties. [22]

## Materials and methods

### Determining the influence of D1 and D2 aerosols on stencil wound healing

The compositions were prepared on the basis of dipyridamole (Medisca, USA) and papaverine (Aldrich/Sigma, USA) in two compositions:

D1 (without ascorbic acid): 1% dipyridamole, 1% papaverine hydrochloride; 0.05% chlorhexidine; 10% polyethylene oxide-400; distilled water up to 100%.

D2 (with ascorbic acid): 1% dipyridamole, 1% papaverine hydrochloride; 0.05% chlorhexidine; 0.25% ascorbic acid; 10% polyethylene oxide-400; distilled water up to 100%.

Both aerosol variants are equivalent in terms of the ratio of active ingredients (1% dipyridamole, 1% papaverine hydrochloride (D1, 0.05% chlorhexidine as a preservative), and for the second version of the aerosol (D2) it is 5% sodium ascorbate). Aerosol solutions were injected into a mechanical nebulizer, and pressurized balloons were not used. To enhance the penetration of active substances into the affected tissues, 10% polyethylene oxide-400 was used as an excipient in both versions of the aerosol [23] Similar solutions were also prepared for intraperitoneal administration, but without chlorhexidine and polyethylene oxide-400. The concentration of ascorbic acid in the D2 variant was reduced to 0.25%.

The wound healing action was studied on the basis of a model of stencil wounds in rats.

Their effect on the number of animals’ own stem cells, CD34, was also evaluated.

The study of the wound healing properties of experimental aerosols D1/D2 was conducted on 38 Wistar male rats.

The experiments involved Wistar male rats weighing 160– 200 g. The animals were obtained from the Biomodelservice (Kiev, Ukraine) and kept in vivarium in Mechnikov Institute of microbiology and immunology. The animals were cared for according to the international guidelines GOST 33647-2015 (Principles of the Good Laboratory Practice, GLP), the international recommendations of the European Convention for the Protection of Vertebrate Animals used for Experimental and Other Scientific Purposes (The European Convention, 1986). The protocol of the experimental study was approved by the Ethics Committee of Mechnikov Institute of Microbiology and Immunology of National Academy of Medical Sciences (protocol № 3-2018 of 14.12.2018).

The animals were divided into four groups: 1 – wounds treated with D1 aerosol (n = 10); 2 – wounds treated with D2 (n = 10); 3 – wounds treated with the reference drug dexpanthenol (Panthenol-Ratiopharm, 5% ointment) (n = 10); 4 – control group (n = 8). Dexpanthenol is a vitamin B5 - well-known substance in the world for tissues regeneration stimulating [24,25]

Stencil wounds [26] of the back were simulated in animals under anesthesia with diethyl ether. The initial wound size was 4 ± 1.0 cm^2^. The application of drugs was initiated from the first day after the alteration. The average amount of aerosol was 0.5 ml/animal. Applications were performed twice a day to make sure that the drug covers the entire surface of the wound and approximately 0.5 cm of depilated skin around it. Then, the animals were fixed for 30 minutes to ensure the absorption of the solution and to prevent them from licking wounds. A planimetric study [26] was conducted on the 3rd, 7th, 10th, 14th and 18th day from the beginning of the experiment. The obtained parameters of the wound surface area were statistically processed using the analysis of variance.

### Stimulating the growth of CD34 + pluripotent hematopoietic cells

The drugs were studied in an experiment on a model of cytostatic hemoimmunosuppression. The experiments were conducted on 90 male Wistar rats weighing 160–200 g. The animals were kept under conditions of free access to food and water. Two experimental and one reference groups consisted of 30 animals each. In both groups, hemoimmunosuppression was caused by a five-fold intraperitoneal administration, at 24-hour intervals, of cyclophosphan (Kyivmedpreparat, Ukraine) in a dose of 10 mg/kg, and prednisolone (Darnytsia, Ukraine) 2 mg/kg in 3 ml of normal saline. On the next day after the completion of the administration of hemoimmunosuppressive drugs, the animals were intraperitoneally injected with a D1/D2 composition in form of IV injection:

D1 (without ascorbic acid): 1% dipyridamole, 1% papaverine hydrochloride; distilled water up to 100%.

D2 (with ascorbic acid): 1% dipyridamole, 1% papaverine hydrochloride; 0.25% ascorbic acid; distilled water up to 100%.

D1/D2 used in the quantity of 0.1 ml/animal (2.5 mg/kg).

At the same time, animals from the reference group were administered, similarly to the experimental group, 0.1 ml of reference normal saline. The procedure was performed daily, once a day for 60 days. Blood samples were taken from the tail vein on days 1, 10, 30 and 60. The proliferative activity of CD34 + stem cells was studied using a CD34 Countkit [27,28] kit for cytofluorimetry via a FACS Calibur flow cytofluorometer. The digital material was processed using the analysis of variance, Student’s/ Bonferroni modification (Fig.1).

**FIG. 1.**
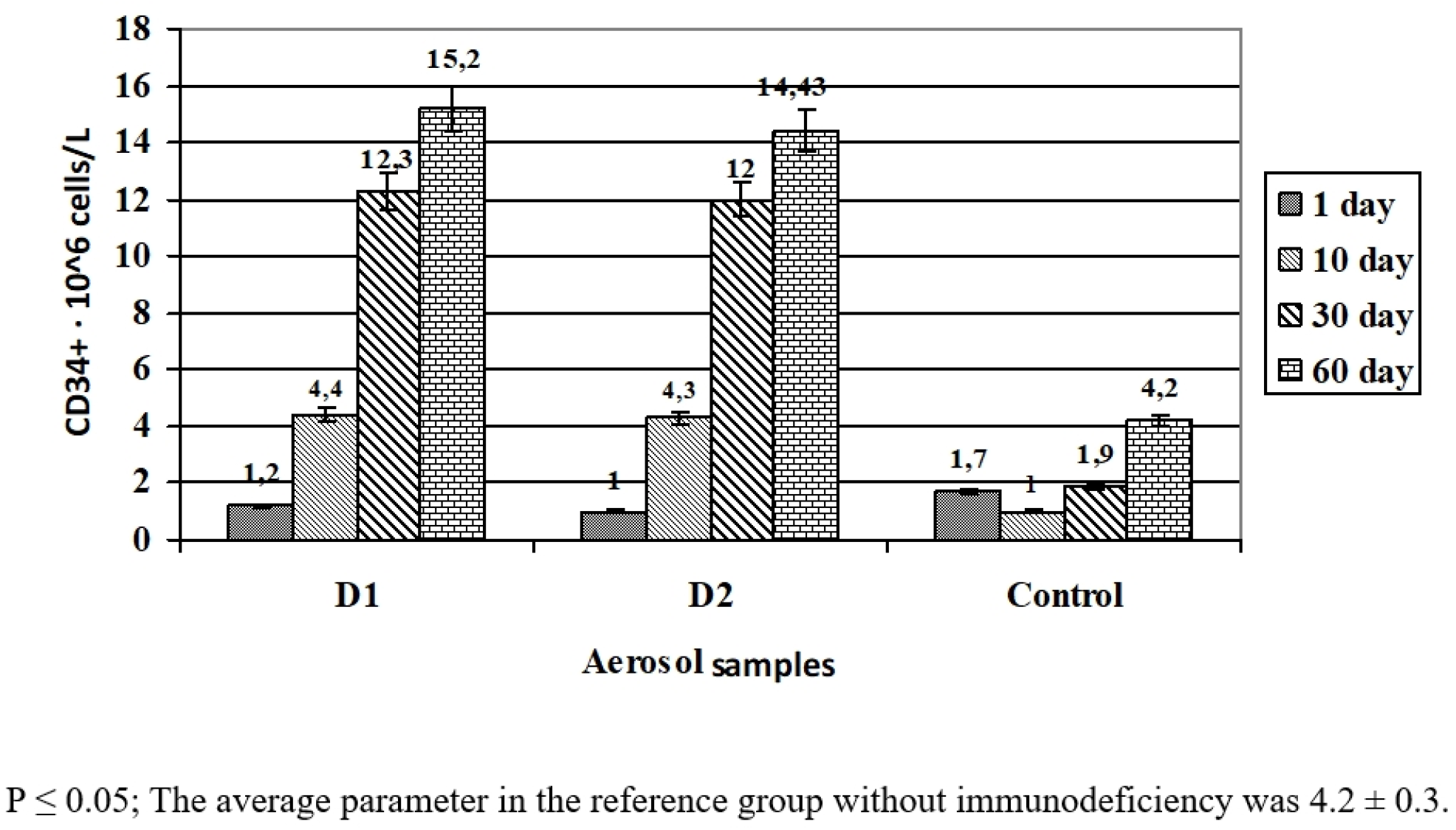
The number of pluripotent CD34 + cells in the blood of rats as affected by the compositions D1 and D2

## Results and discussion

The aerosol of the two compositions had a significant impact on the dynamics of the wound process in rats. On day 3 (48 hours after the simulation of stencil wounds), the maximum intensity of the inflammatory process was noted in the pathology control group; there were no statistical differences between the experimental and the reference groups. Oedema and hyperemia affected the surrounding tissues as well. During this period, the area of the wound surface even slightly exceeded the initial parameters (Table 1). The general condition of the animals was severe. They were sedentary, their coat was disheveled, and they ate less food. During this period, the experimental aerosols D1 and D2 did not cause any changes in the area of wounds; however, less pronounced local inflammatory reaction and a somewhat better general condition of animals were noted.

**TABLE. 1.** The effect of D1/D2 aerosols on the surface area of stencil wounds in rats

**TABLE. The effect of D1/D2 aerosols on the surface area of stencil wounds in rats**
| Drug product | n | Wound surface area, cm<sup>2</sup> (M±m) |  |  |  |  |
| --- | --- | --- | --- | --- | --- | --- |
|  |  | Day 3 | Day 7 | Day 10 | Day 14 | Day 18 |
| D1 | 10 | 4.36**±0.74 | 3.31*<sup>1</sup>±0.71 | 2.58*<sup>1</sup>±0.79 | 0.72*<sup>1</sup>±0.23 | - |
| D2 | 10 | 4.21**±1.06 | 3.40*<sup>1</sup>±0.87 | 2.12*<sup>1</sup>±0.73 | 0.78*<sup>1</sup>±0.70 | - |
| Dexpanthenol | 10 | 4.47**±0.36 | 4.29**±0.30 | 3.98**±0.44 | 1.66 *±0.32 | - |
| Wound control | 8 | 4.18±0.56 | 4.04±0.50 | 3.88±0.52 | 2.55±0.38 | 0.62±0.22 |

\* $P \leq 0.05$ (dispersions differ significantly versus the wound control parameters for the corresponding period)
\*\* $P > 0.1$ (dispersions do not differ versus the wound control parameters for the corresponding period)
Starting from day 7, the differences between D1/D2 and the dexpanthenol group were statistically significant at $P \leq 0.05$

Subsequently, the experimental groups showed a fast positive dynamics in the clinical condition of rats. On day 7, hyperemia and oedema in the experimental groups were insignificant, and granulation tissue was formed in the center of the wound. The onset of reparative processes was noted: marginal occlusion of the defect, which changed the wound area. During this period, a significant decrease in the area of the wound surface was observed vis-à-vis the parameters of day 3. At the same time, this parameter did not change in the pathology control group.

On the 10th day of the experiment, marginal epithelialization of wounds, a significant decrease in the area of the wound surface in comparison to the baseline data and untreated control, complete cleansing of the wound surface, the presence of granulations, and the absence of inflammation were noted in both experimental groups.

The subsequent periods saw a rapid healing of the wound defect in rats treated with D1/D2 aerosols. On the 14th day, the wounds were dry and their area was 0.72 and 0.78 in groups 1 and 2, respectively, which meant that most of the wound surface was epithelialized. At the same time, although a number of animals in the pathology control group showed positive dynamics, four out of eight rats still had an inflammatory process, and the area of the wound surface was large.

Statistically significant differences between the experimental groups (including dexpanthenol group) and the reference group were noted as early as on day 7. During the same period, as well as on days 10 and 14, significant advantages of the experimental sprays vis-à-vis the action of dexpanthenol were observed. By day 18, wounds were completely healed in all rats of the experimental groups, while in the reference group, three animals did not complete wound epithelialization and one of them had pus discharge.

There were no statistically significant differences between the D1 and D2 groups. Therefore, ascorbic acid had no pronounced influence on the wound healing effect produced by the spray with dipyridamole and papaverine.

Dexpanthenol ointment also helped reduce the wound healing time. However, its effect was somewhat less pronounced than the effect from the use of aerosols.

The figure shows the results of studying the effect of the compositions on the number of pluripotent CD34 + cells in the blood of rats. The normal level of CD34 cells in the blood ranges from 3 to 6 * 10 ^6^ cells/liter.

The administration of cytostatic drugs led to a three-to fourfold decrease in the level of CD34 by the time of the start of therapy comparing to the baseline level. Dipyridamole with papaverine in both compositions had a stimulating effect on the processes of bone marrow regeneration and restoration of peripheral blood parameters, increasing the level of CD34 to normal values. By day 30, its level increased more than twofold thanks to the effect from D1 and D2 samples, while in the animals from the reference group, it returned to normal values only on the 60th day. Therefore, dipyridamole with papaverine caused about a twofold acceleration of these processes.

The maximum effect from the use of the compositions was observed on the 60th day, and the restoration of the physiological level of pluripotent cells was observed even on the 10th day after the start of their use. The increase in the number of pluripotent cells coincided in time with the stimulation of reparative processes (on the model of stencil wounds in rats), which apparently indicates the predominant stimulation of stem cell division at the periphery rather than the enhancement of bone marrow functions.

The presence of ascorbic acid in the compositions had no bearing upon their effect on stimulation of the production of CD34 cells.

## Conclusions

1. Dipyridamole- and papaverine-based aerosols of two compositions (with and without ascorbic acid) have pronounced reparative properties, significantly accelerating epithelialization and healing of stencil wounds in rats. In terms of this type of action, they are somewhat superior to dexpanthenol.
2. Dipyridamole- and papaverine-based aerosols have the ability to produce beneficial effect on the entire body’s immune system by stimulating the division of pluripotent CD34 cells.
3. The combined effect of papaverine and dipyridamole on tissues leads to selective stimulation of the division of pluripotent cells in the wound, and contributes to a sixfold acceleration of restoration of the animal’s immune system after induced immunodeficiency.
4. The absence of a statistically significant difference between the groups of D1 and D2 compositions indicates an insignificant role played in wound regeneration by ascorbic acid in the aforementioned dosage.

## References

[1] Lipsky BA et al. Diagnosis and treatment of diabetic foot infections. Clinical Infectious Diseases 2004; 39(7): 885–910.

[2] Lee KB et al. Topical embryonic stem cells enhance wound healing in diabetic rats. J Orthop Res 2011; 29: 1554–1562.

[3] Howe AK. Regulation of actin-based cell migration by cAMP/PKA. Biochim Biophys Acta 2004; 1692: 159–174.

[4] Laird DJ et al. Stem cell trafficking in tissue development, growth, and disease. Cell 2008; 132: 612–630.

[5] Saidy K, Al-Alaiyan S. The use of F-arginine and phosphodiesterase inhibitor (dipyridamole) to wean from inhaled nitric oxide. Indian J. Pediatr 2001; 68(2): 175–177.

[6] Galabov AS, Mastikova M. Dipyridamole induces interferon in man. Biomed. Pharmacother. Biomédecine pharmacothérapie 1984; 38(8); 412–413

[7] Triner L et al. Cyclic phosphodiesterase activity and the action of papaverine. Biochemical and biophysical research communications 1970; 40(1): 64–69.

[8] Telang P. Vitamin C in dermatology. Indian Dermatol Online J 2013; 4(2):143–6.

[9] Lima CC, Pereira AP, Silva JR, Oliveira LS, Resck M, Grechi CO, et al. Ascorbic acid for the healing of skin wounds in rats. Braz J Biol 2009; 69(4):1195–201.

[10] Ghosh O. et al. Phosphodiesterase inhibitors: their role and implications. Int J PharmTech Res 2009; 1 (4): 1148–1160

[11] Lo KW et al. One-day treatment of small molecule 8-bromo-cyclic AMP analogue induces cell-based VEGF production for in vitro angiogenesis and osteoblastic differentiation. J Tissue Eng Regen Med 2013; DOI:10.1002/term/1839.

[12] Rodriguez-Pallares J. et al. Dipyridamole-induced increase in production of rat dopaminergic neurons from mesencephalic precursors. Neurosci. Lett 2002; 320 (1-2): 65–68

[13] Tornling G et al. Proliferative activity of myocardial capillary wall cells in dipyridamole-treated rats. Сardiovascular research 1978; 12: 692–695

[14] Snoeck R et al. Dipyridamole potentiates the activity of various acyclic nucleoside phosphonates against varicella-zoster virus, herpes simplex virus and human cytomegalovirus. Antivir Chem Chemother 1994; 5(5):312–321

[15] Tenser RB., Gaydos A, Hay K A. Inhibition of herpes simplex virus reactivation by dipyridamole. Antimicrob Agents Chemother 2001; 45(12): 3657–3659

[16] Galabov AS, Mastikova M. Interferon-inducing activity of dipyridamole in mice. Acta Virol 1983; 27(4): 356–358

[17] Zhilinskaya IN et al. Antiviral activity of some vasodilative preparations. Bull Exp Biol Med 1996; 121(2): 154–156

[18] Perez RM. Antiviral activity of compounds isolated from plants. Phar. Biol 2003; 41(2): 107–157

[19] Turano A. et al. Inhibitory effect of papaverine on HIV replication in vitro. AIDS Res Hum Retroviruses 1989; 5(2): 183–192

[20] Basceken SI. et al. Effects of papaverine on healing of colonic anastomosis in rats. European Surgery 2017; 49(4): 158–164.

[21] Kuwano H. et al. Dipyridamole inhibits early wound healing in rat skin incisions. Journal of Surgical Research 1994; 56(3): 267–270

[22] Sanchez IR. et al. Effects of Chlorhexidine Diacetate and Povidone‐lodine on Wound Healing in Dogs. Veterinary surgery 1988; 17(6): С. 291–295

[23] Khokhlenkova NV. Prospects of Using Biopolymeric Films in Medicine and Pharmacy. Asian Journal of Pharmaceutics 2017; 11(04): 672–677.

[24] Semieka MA. et al. Comparative Study of the Therapeutic Effect of Panthenol Gel and Mebo Ointment on Metacarpal Wound Healing in Donkeys. Journal of Equine Veterinary Science 2019; 74:21–27

[25] Proksch, E. et al. Topical use of dexpanthenol: a 70th anniversary article. Journal of Dermatological Treatment 2017; 28(8): 766–773.

[26] Cheraghali Z. et al. Planimetric and biomechanical study of local effect of pulegone on full thickness wound healing in rat. The Malaysian journal of medical sciences 2017; 24(5): 52

[27] Sutherland DR, Keeney M. Enumeration of CD34+ cells by Flow Cytometry //Cellular Therapy: Principles, Methods and Regulations. An American Association of Blood Bankers Cell Therapy Technical Manual. – The American Association of Blood Bankers (AABB), Bethesda MD, 2009. P. 538–554.

[28] Murugesan M. et al. Flow cytometric enumeration of CD34+ hematopoietic stem cells: A comparison between single-versus dual-platform methodology using the International Society of Hematotherapy and Graft Engineering protocol. Asian Journal of Transfusion Science 2019; 13(1): 43–44.

